# Precision Functional Mapping of the Individual Human Brain Near Birth

**DOI:** 10.1101/2025.07.07.663543

**Authors:** Alyssa K. Labonte, Julia Moser, M. Catalina Camacho, Jiaxin Cindy Tu, Muriah Wheelock, Timothy O. Laumann, Evan M. Gordon, Damien A. Fair, Chad M. Sylvester

**Author notes:** Corresponding Author: Department of Psychiatry, Washington University, 660 S. Euclid, Campus Box 8511, St. Louis, MO 63110, USA.

## Abstract

Cortical areas are a fundamental organizational property of the brain, but their development in humans is not well understood. Key unanswered questions include whether cortical areas are fully established near birth, the extent of individual variation in the arrangement of cortical areas, and whether any such individual variation in cortical area location is greater in later-developing association areas as compared to earlier-developing sensorimotor areas. To address these questions, we used functional MRI to collect precision functional mapping (PFM) data in eight individual neonates (mean 42.7 weeks postmenstrual age) over 2-5 days (mean 77.9 minutes of low motion data per subject [framewise displacement <0.1]). Each subject’s dataset was split into two roughly equal halves of data from different days of data collection to measure within-subject reliability and across-subject similarity. Whole-brain patterns of functional connectivity (FC) reached a mean within-subject, across-day reliability of r=0.78 with 41.9 minutes of data. Across subject similarity of whole-brain FC was r=0.62 on average and significantly lower than within-subject similarity (t=5.9, p<0.001). Using established methods to identify transitions in FC across the cortical surface, we identified sets of cortical areas for each individual that were subject-specific and highly reliable across split-halves (mean z=4.4, SD=1.4). The arrangement of cortical areas was thus individually specific across the entire cortical surface, and this individual specificity did not vary as a function of the sensorimotor-association axis. This study establishes the feasibility of neonatal PFM and suggests that cortical area arrangement is individually specific and largely established shortly following birth.

**GRAPHICAL ABSTRACT:** 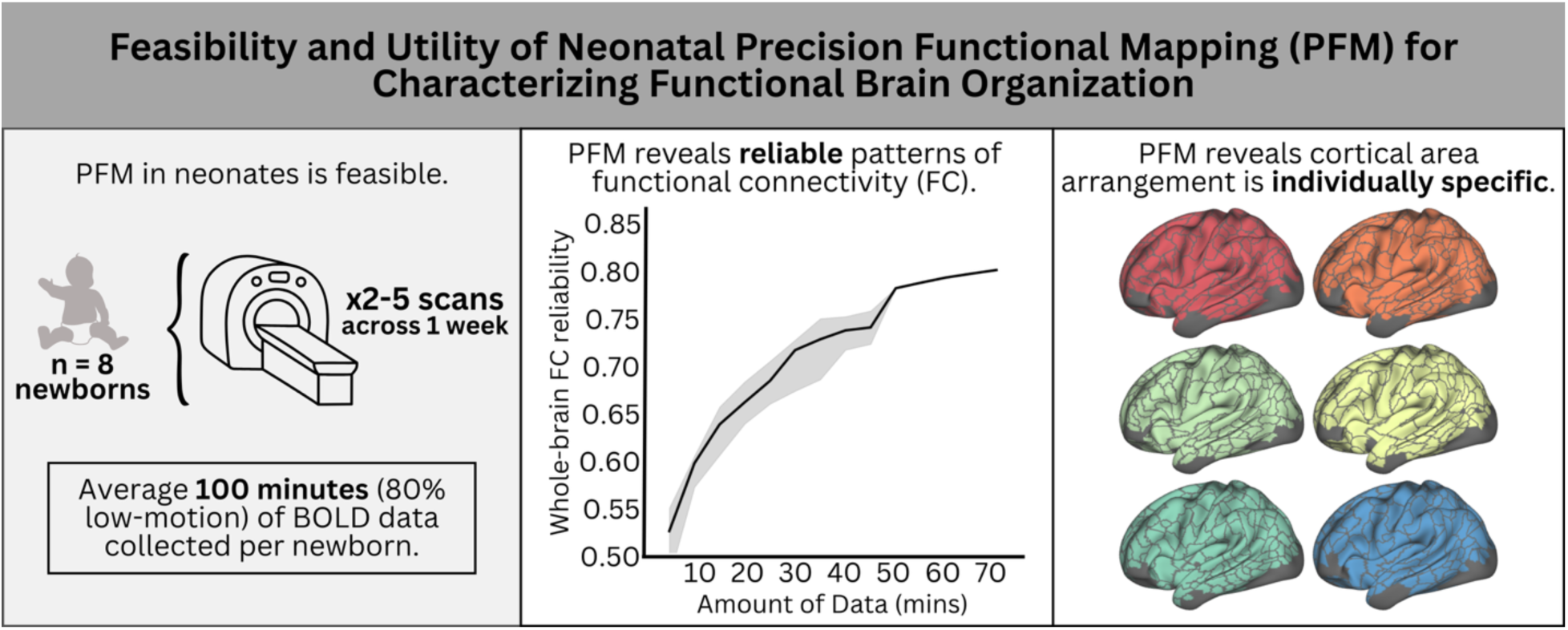

**HIGHLIGHTS:**

- It is feasible to obtain precision functional mapping (PFM) in neonates, acquiring 60+ minutes of fMRI data in individual neonates over multiple days.
- Neonatal functional connectivity (FC) measures obtained through PFM are moderately reliable with 40 minutes of data.
- Patterns of neonatal FC across the brain are individually specific.
- Cortical areas, based on patterns of homogenous FC, can be reliably identified across the entire cortical surface in individual neonates.
- The arrangement of cortical areas at birth is individually specific across the entire brain.

## INTRODUCTION

Individual variation in functional brain organization relates to individual differences in behaviors and psychiatric risk across the lifespan (Finn et al., 2015). This individual variation likely emerges through development (Fox et al., 2015; Pérez-Edgar et al., 2020), but when and how individual differences in functional brain organization emerge is poorly understood. This knowledge gap impedes our ability to create developmental models of individual differences that result in subsequent variation in behavior and psychiatric risk. Precision functional mapping (PFM) has been used in human children and adults to map individual functional brain organization (Gordon et al., 2017b, 2023; Laumann et al., 2015). However, its feasibility and utility in neonates has not been clearly demonstrated. This study uses PFM in individual neonates to test the hypothesis that individual variation in a fundamental organizational property of the human brain – the arrangement of cortical areas across the cortical surface – is already present near the time of birth.

The division of the human cerebral cortex into distinct cortical areas is a key component of the brain’s hierarchical organization and central to higher-order cognition and behavior (Petersen et al., 2024). Adjacent cortical areas can be distinguished from one another based on differential patterns of connectivity to other regions of the brain (Felleman and Van Essen, 1991; Petersen et al., 2024; Sejnowski and Churchland, 1989). The putative locations and arrangement of cortical areas in humans can thus be identified by using functional MRI (fMRI) to identify contiguous portions of cortex with homogeneous patterns of functional connectivity (FC) that are distinct from neighboring cortex, an approach known as the boundary mapping (Cohen et al., 2008; Glasser et al., 2016; Gordon et al., 2016; Schaefer et al., 2018; Wig et al., 2014). Work using this technique has revealed a shared topology in the arrangement of cortical areas across adults and children as young as 2 years old (Gordon et al., 2016; Tu et al., 2025a). Despite these similarities across individuals of different ages, additional work has noted individual variation in the exact placement and size of different cortical areas (Gordon et al., 2017b; Laumann et al., 2015), which may relate to individual differences in behavior, emotions, and risk for brain-based illnesses (Lynch et al., 2024; Seitzman et al., 2019).

The division of the cortex into cortical areas emerges during early development, a process known as arealization (O’Leary et al., 2007). Inter-individual variation in the details of arealization during development could account for variation in the ultimate arrangement of cortical areas in adulthood and their associated behavioral phenotypes. Yet, little is known about the development of cortical areas in humans, including whether cortical areas are fully established near birth. Animal models have demonstrated that discrete cortical areas develop from a continuous “protomap” established during embryonic development, with refinement of areal borders and discretization of areal anatomy and function driven by postnatal sensory input through thalamocortical afferents (Cadwell et al., 2019; O’Leary et al., 2007). Thus, postnatal experience may be necessary for the complete development of discrete cortical areas.

Another aspect of development that remains unknown is whether the maturation of cortical areas is uniform across the cortex; or if sensory cortical areas mature prior to association areas. Maturation of brain-wide FC occurs asynchronously across the cortex during the early postnatal period along a sensorimotor-association (S-A) axis (Gao et al., 2009, 2015b), whereby FC of sensorimotor regions matures prior to FC of association regions (Doria et al., 2010; Gao et al., 2015b, 2015a, 2009; Smyser et al., 2010). This model of S-A axis development (Sydnor et al., 2023) also parallels postnatal experience, in which primary somatosensory inputs precede more complex environmental inputs (Du et al., 2024). One possibility is that the development of cortical areas mirrors this same developmental trajectory, such that postnatal experience drives maturation of areal boundaries in sensory areas prior to association areas. In this cascading model, the borders of primary sensory areas are first refined by initial sensory inputs both pre- and postnatally. Secondary areas then receive inputs from previously refined primary sensory areas and thus develop after the primary areas are refined. Similarly, multimodal association cortex areas only start developing once the secondary areas are refined, following the same developmental hierarchy.

In prior work, we used the boundary mapping technique to identify the arrangement of cortical areas shortly after birth, using fMRI data averaged across 131 neonates (Myers et al., 2024). In contrast to adults and children, in neonates, we found it was necessary to shrink area parcels to their centers to obtain a reliable fit to held out data. We reasoned that this observation has at least two possible explanations: 1) cortical arealization remains immature in the early postnatal period such that area boundaries may not be firm; or, 2) individual variation in the arrangement of cortical areas across the brain in neonates blurs cortical boundaries when averaged. One method to adjudicate between these explanations would be to define cortical areas in individual neonates as opposed to group-average data.

Precision functional mapping (PFM), in which large amounts of fMRI data are collected in individuals across multiple days, allow for precise and individualized characterization of functional brain connections, cortical areas, and networks (Gordon et al., 2017b, 2017a; Laumann et al., 2015). PFM has uncovered individual-specific features of functional brain organization, such as shifts in areal boundaries obscured by group-level analyses, that are crucial for advancing our understanding of brain-behavior relationships and cortical organization (Gordon et al., 2023, 2017b; Gordon and Nelson, 2021; Gratton et al., 2020; Michon et al., 2022; Seitzman et al., 2019). Group-averaging may blur individual variability to an even greater extent in neonates, given that rapid neurodevelopment during this epoch may mean that not all individuals are at the same developmental stage. Thus, individual approaches are essential for understanding these earliest stages of postnatal brain organization. However, little is known about using PFM in neonates, including whether the technique is feasible in this age range, how much data is required in neonates to have individually reliable measures of FC, or even whether the brain organization of different neonates is unique enough to warrant PFM studies.

Here, we use PFM to characterize FC reliability and functional brain organization near birth in 8 individual neonates with between 40.2 and 154.1 minutes of data in each individual. We hypothesized that, as in older populations, it would be possible to identify putative cortical areas across the entire cortex in each neonate that are reliable and individual-specific in their arrangement. Further, because FC is thought to develop along a S-A axis, we hypothesized that the degree of individual specificity in cortical area arrangement would be greater in higher-order association areas relative to sensorimotor areas. Overall, this study provides insights into the feasibility and reliability of neonatal PFM as well as key insights into the state of cortical areas near birth.

## MATERIALS AND METHODS

This study was approved by the Human Research Protection Office at Washington University in St. Louis. All parents/guardians of neonatal participants provided informed consent prior to study initiation. The primary dataset in this study, Precision Baby (PB), collected fMRI data from eight healthy, full-term neonates (average postmenstrual age (PMA) 42.7 weeks, range 40-44 weeks; **Table 1**) each scanned across 2-5 consecutive days with the goal of collecting at least 60-minutes of low-motion fMRI data (framewise displacement <0.1-mm) in each individual. This data set was collected for a variety of scientific goals, including piloting for the HEALthy Brain and Child Development (HBCD) study (Dean et al., 2024); thus, protocols and data collection goals differed across participants. Recruitment of the PB cohort and scanning took place between October 2021 and September 2023.

**Table 1.** Participant Demographics.

| PB ID | PMA at Scan 1<br>(weeks) | Sex | Race | Ethnicity |
| --- | --- | --- | --- | --- |
| PB001 | 42.0 | Female | African American or Black | Non-Hispanic |
| PB003 | 43.3 | Male | White | Non-Hispanic |
| PB004 | 44.4 | Female | African American or Black | Non-Hispanic |
| PB006 | 41.7 | Female | White | Non-Hispanic |
| PB008 | 42.1 | Male | White | Non-Hispanic |
| PB012 | 40.9 | Male | White, Asian | Non-Hispanic |
| PB013 | 43.6 | Male | White | Non-Hispanic |
| PB014 | 43.0 | Male | White | Non-Hispanic |

Neuroimaging was performed during natural sleep using a Siemens 3T PRISMA scanner and 32-channel infant specific head coil. Prior to scanning, neonates were fed, swaddled, and positioned in a head-stabilizing vacuum fix wrap (Mathur et al., 2008). Both a T2-weighted (0.8- mm isotropic resolution, time to echo (TE)=563ms, repetition time (TR)=4500ms) and T1- weighted (0.8-mm isotropic resolution, TE=2.22ms, TR=2400ms) image were collected in each neonate. Functional images and spin-echo field maps were collected using various parameters across each of the individual neonates (**Supplementary Table 1**). Framewise integrated real-time MRI monitoring (FIRMM) was used during scanning to monitor real-time neonate movement (Badke D’Andrea et al., 2022; Dosenbach et al., 2017).

### fMRI Processing

All individuals in the PB dataset underwent identical preprocessing and FC processing using in-house pipelines with the exception of PB001 which was collected using multi-echo (ME) sequences. For this subject, motion regressors were calculated on the first echo and applied to all individual echos before they were optimally combined using a T2* based echo weighting approach. BOLD preprocessing steps were equivalent to the rest of the PB subjects after optimal echo combination. BOLD preprocessing included correction of intensity differences attributable to interleaved acquisition, linear realignment within and across runs to compensate for rigid body motion, bias field correction, intensity normalization of each run to a whole-brain mode value of 1000, and distortion correction in native volumetric space. Field distortion correction was performed with the FSL TOPUP toolbox (http://fsl.fmrib.ox.ac.uk/fsl/fslwiki/TOPUP) using the single best field map across all runs for each scanning session based on image quality. BOLD images were then linearly registered to the adult MNI152 atlas (Mazziotta et al., 2001a, 2001b, 1995) as follows: BOLD → individual T2-weighted image → cohort-specific T2-weighted atlas → 711-2N Talairach atlas (adult space) → MNI152 atlas (adult space). The cohort-specific T2 atlas was created from a subset of 50 neonates from the Early Life Adversity and Biological Embedding (eLABE) dataset (Lean et al., 2022; Myers et al., 2024; Nielsen et al., 2022; Sylvester et al., 2022). The volumetric preprocessed BOLD data were then mapped (ribbon-constrained) to subject-specific surfaces prior to FC processing. The Melbourne Children’s Regional Brain Atlases (MCRIB) (Adamson et al., 2020; Alexander et al., 2017), an approach to surface-based neonatal tissue segmentation, was used to generate surfaces for each subject from their T2 image in native space. Subject-specific surfaces then underwent linear transformation to adult MNI152 space and were finally aligned into the “fsLR_32k” surface space using spherical registration procedures (Glasser et al., 2013) adapted from the Human Connectome Project as implemented in Connectome Workbench 1.2.3 (Marcus et al., 2013, 2011). All volumetric and surface registrations were visually inspected to ensure quality and accuracy.

Following initial volumetric preprocessing and surface registration to fsLR_32k space with a small smoothing kernel (σ =1-mm), BOLD time series were censored at framewise displacement (FD) <0.1 mm, and only epochs of at least 3 consecutive frames with FD <0.1 mm were included (**Supplementary Figure 1**). Runs with <50% of frames retained were excluded. Neonates in the PB dataset with less than 60-minutes of data remaining after frame censoring were excluded from all split-half analyses (n=2 excluded from split-half analyses).

Functional data underwent FC processing as follows (Power et al., 2014): (i) demean and detrend within each run, ignoring censored frames; (ii) multiple regression with nuisance time series including white matter, ventricles, and whole brain (average gray matter signal), as well as 24 parameters derived from head motion, ignoring censored frames. Finally, retained data were interpolated at censored timepoints to allow band-pass filtering (0.005 Hz < f <0.1 Hz). Time courses for surface data were smoothed with geodesic 2D Gaussian kernels (σ=2.25-mm) after FC processing and concatenated across all sessions within a single individual, excluding censored frames. FC was computed as the Fisher z-transformed Pearson correlation between time courses from pairs of surface vertices.

### Functional Connectivity Reliability and Similarity

Following methods outlined in (Gordon et al., 2017b; Laumann et al., 2015), we evaluated the reliability and similarity of FC patterns within- and across-subjects. To evaluate the reliability of FC, each subject’s dataset was split into two approximately equal-sized subsets (**Supplementary Table 2**). Split-half datasets were created by combining data across scan sessions to maximize data quantity (post motion censoring) in each split-half. Data within a single scan session were not broken up between each split-half of data, but combined scan sessions were not necessarily temporally adjacent. For all subjects, the maximum amount of held-out data was selected from the split-half with the most data. Then, a varying amount of continuous data was selected from the second split-half in increments of 5-minutes and the time courses for each split-half were correlated against one another to generate an FC matrix for each split-half of data. Finally, vertex-wise FC matrices for each split-half were compared by correlating the upper triangles of the matrices to obtain a single Pearson’s r value. This procedure was repeated 1000 times with a different random selection (in continuous 5-minute increments) of data in each iteration.

To evaluate the similarity of FC patterns within- and across-subjects, we calculated the pairwise similarity between all individual subject’s split-half datasets. First, we generated individual vertex-wise FC matrices for each split-half dataset for each subject, as described in fMRI processing. Then, we calculated a ‘similarity matrix’ by correlating the upper triangle of each split-half dataset’s FC matrix against the upper triangle of all other split-half datasets across all subjects. We also computed the ‘similarity matrix’ between each split-half dataset from each subject and a group average FC matrix. For this latter computation, the individual’s own split-half datasets were excluded from the average FC matrix to minimize bias.

### Parcel Generation

Previously described boundary mapping methods were used to identify abrupt transitions in FC across the cortical surface (Gordon et al., 2016; Myers et al., 2024) and are described briefly here. First, whole-brain FC maps at each cortical surface vertex were computed. Then, FC map similarity between all vertices was computed. Spatial gradients were calculated on each column of this similarity matrix using Workbench tools (wb_command -cifti-gradient), creating a gradient map for each surface vertex. The resulting gradient maps were smoothed with a kernel of σ=2.55 mm. Next, edges were identified in each smoothed gradient map using the watershed edge detection algorithm (Beucher & Lantuejoul, 1979) whereby gradient maps are “filled up” from their local minima until they reach a peak in the spatial gradient (i.e., rapid changes in FC similarity). These edge maps are averaged across all vertices, creating an edge density map. Parcels were identified by applying the watershed edge detection algorithm to the edge density map, again filling edges from their local minima to a predefined height threshold (here, the 90th percentile of edge density values). A flowchart of this method can be seen in **Supplementary Figure 2**.

As a final step, neighboring parcels with low edge density between them were merged based on a predefined threshold (here, 70^th^ percentile of edge density values). Parcels with at least 15 vertices inside low signal areas (mean BOLD signal <750 after mode 1,000 normalization; see Supplementary Methods and **Supplementary Figure 3**) were excluded (Gordon et al., 2016; Myers et al., 2024; Ojemann et al., 1997).

### Parcel Evaluation

#### Cortical Area ‘Fit’ – FC Homogeneity

Parcels were generated based on abrupt changes in FC patterns across the cortical surface (Gordon et al., 2016). Thus, resulting parcels should be distinct from neighboring parcels in connectivity pattern, as well as show highly homogenous connectivity within itself. To measure the homogeneity of a parcel, a principal components analysis (PCA) was run on the connectivity patterns from all individual vertices comprising a single parcel, and homogeneity was defined as the percent of variance explained by the first principal component (Gordon et al., 2016; Myers et al., 2024). A null distribution of FC homogeneity from 1000 randomly rotated parcellations having parcels of the same size, shape, and configuration as the true parcellation was computed for each individual. The average homogeneity across all parcels for each null parcellation was then compared to the average homogeneity of the true parcellation, calculated as a z-score. This procedure was done using the method originally described in (Gordon et al., 2016).

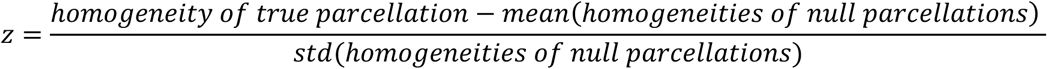

#### Individual Specific Cortical Area Reliability and Similarity

To test the within-subject reliability and across-subject similarity of each subject’s cortical area parcellation, we split each subject’s dataset into two halves (split-half 1 and split-half 2; described above), generated parcellations from each split-half independently, and evaluated the fit and spatial overlap of the resulting cortical area parcels and their boundaries (see *Cortical Area ‘Fit’ – FC Homogeneity* and Supplementary Methods). For all across-subject similarity analyses, split-half 1 cortical parcels for each neonate were tested in data from split-half 2 of every other neonate’s dataset.

### Regional Specificity

To examine whether the locations of cortical areas might be consistent across individuals in specific parts of the brain, we generated a whole-brain map representing the average homogeneity of cortical areas. This map was generated by assigning the average homogeneity value calculated from across-subject cortical area similarity comparisons (described above) for each vertex across the entire cortical surface. A Pearson correlation coefficient was computed between this map of average homogeneity and the sensorimotor-association axis (S-A axis) (Sydnor et al., 2023). We also computed a null distribution of 1000 randomly rotated S-A axis schemes for significance testing. The average homogeneity was correlated with each null S-A axis scheme and was then compared to the average homogeneity of the true S-A axis scheme, calculated as a z-score.

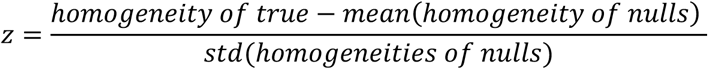

Additionally, region-specific homogeneity was computed as the average within-region homogeneity for sensorimotor and association regions along the sensorimotor-association axis (S-A axis) (Sydnor et al., 2023). For this calculation, the S-A axis map was binarized whereby vertices with values greater than 25000, were classified as sensorimotor regions and those less than 25000 were classified as association regions. We computed a null distribution of 1000 randomly rotated binary S-A axis schemes for significance testing. The average homogeneity across all vertices for each null S-A axis scheme was then compared to the average homogeneity of the true network scheme, calculated as a z-score (see equation above).

## RESULTS

### Neonatal precision neuroimaging is feasible and is moderately reliable with 40 minutes of data

We collected FC data in eight neonates within the first several weeks of birth. **Table 1** shows demographic information for this Precision Baby (PB) dataset, and **Figure 1A** lists the amount of data collected and retained for each neonate following motion censoring procedures best suited to this dataset (FD<0.1mm; **Supplementary Figure 1**). On average, we obtained 99.5 minutes of scan data per neonate (range 40.2–154.1 mins) over 2–5 consecutive days, with an average of 77.9 minutes of data (range 36.6–143.8 mins) remaining post-motion censoring. Only neonates with at least 60-minutes of low-motion data were included in remaining analyses (n=6). Measures of FC were moderately reliable within-subject across multiple scanning sessions (**Figure 1B**). The average within-subject reliability was r=0.78 (SD=0.03). Note that this analysis requires half of each subject’s data to be held out and equal and thus assessed reliability including an average of 41.9 minutes of data (range 29.5–69.8). Results were similar when matching total amounts of data included for each subject in the held-out half (31.7 minutes of data (**Supplementary Figure 4**)). Reliability increased as a function of total amount of data included, with no clear plateau in reliability even with the maximum amount of data in each subject. This result suggests that further reliability increases are expected when collecting more than 50-70 minutes of data.

**Figure 1.**
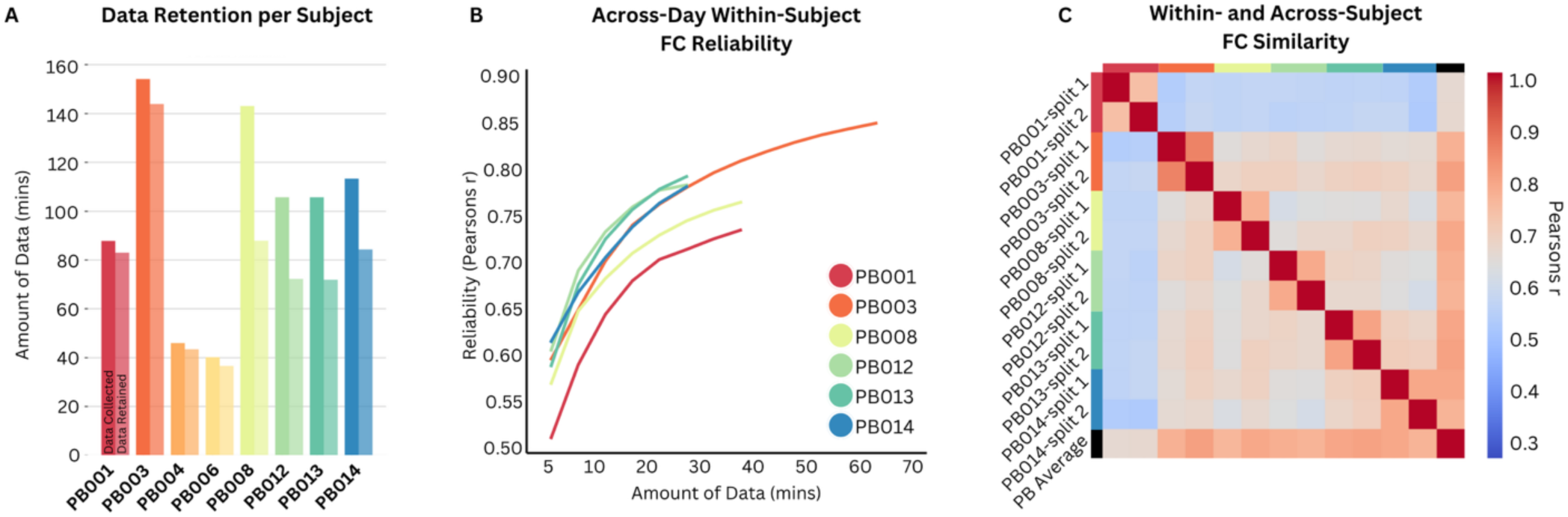
Brain-wide patterns of FC are reliable and individual-specific at birth. **A)** The total amount of data, in minutes, collected (left bar) and retained (right bar) in each participant’s full dataset. **B)** Across-day reliability of whole-brain FC increases with data quantity. Reliability was measured as follows: For each participant, the upper triangle of the FC matrix of a given amount of randomly selected data was correlated with the upper triangle of the FC matrix from the split-half of their data with the most retained data (see Supplementary Table 2) to obtain a Pearsons r correlation for that particular amount of data. This procedure was repeated 1000 times for each participant. **C)** Pairwise similarity of FC matrices between all individual split-halves of data across all participants, as well as the group average (black; last row and column). Similarity across split-halves within each participant was higher than across participants (t=5.9, p<0.001). Similarity was measured as follows: For each participant, the upper triangle of the FC matrix for each split-half was correlated with the upper triangle of every other split-half FC matrix across all participants, as well as the group average FC matrix.

### Neonatal functional connectivity is individually specific

Measures of FC were less similar across-subjects, suggesting individual-specific patterns of FC beginning at birth. Within vs. between individual FC similarity was evaluated by correlating FC matrices generated from split-halves of every subject’s dataset with every other subject’s split- half dataset. As before, measures of FC were highly similar within individuals across two separate halves of their dataset (mean r=0.78; **Figure 1C**). However, FC similarity across individuals was comparatively low (mean r=0.62, SD=0.06; **Figure 1C**). Across all split-halves, FC similarity was greater within subject than across subjects (t=5.9, p<0.001).

### FC gradient-based cortical areas can be reliably defined in individual neonates and cover ∼90% of the cortical surface

We used a boundary mapping approach to parcellate the neonatal cortex into putative cortical areas tiling ∼90% of the cortical surface (**Figure 2**). Each individual neonate had, on average, 461 cortical areas tiling both hemispheres (range 430-529). The reliability of putative cortical area parcellations was evaluated using split-half data from each individual neonate. The colored dots in **Figure 3** depict homogeneity, a standard metric to assess goodness of fit (Gordon et al., 2016), for cortical parcellations for each individual neonate when tested against their own held-out data (smaller black dots represent the homogeneity of spatially permuted null parcellations, see methods). Across all neonates, cortical area parcellations generated from split-half 1 had homogeneity z-scores significantly greater than chance when tested against split-half 2 (mean z=4.4, all p<0.01), indicating that individual-specific cortical area parcellations are reliable across independent data from the same individual. This result was confirmed using 3 additional metrics assessing the within-subject reliability of the parcels or the boundaries between parcels, including the distance-controlled boundary coefficient and spatial overlap of both parcels and parcel boundaries (see **Supplementary Results** and **Supplementary Figures 5** and **6**). Notably, reliable parcels covered ∼90% of the cortical surface, consistent with results in older children and adults (Gordon et al., 2016; Tu et al., 2025a), but contrary to prior findings in group-averaged neonatal data (Myers et al., 2024).

**Figure 2.**
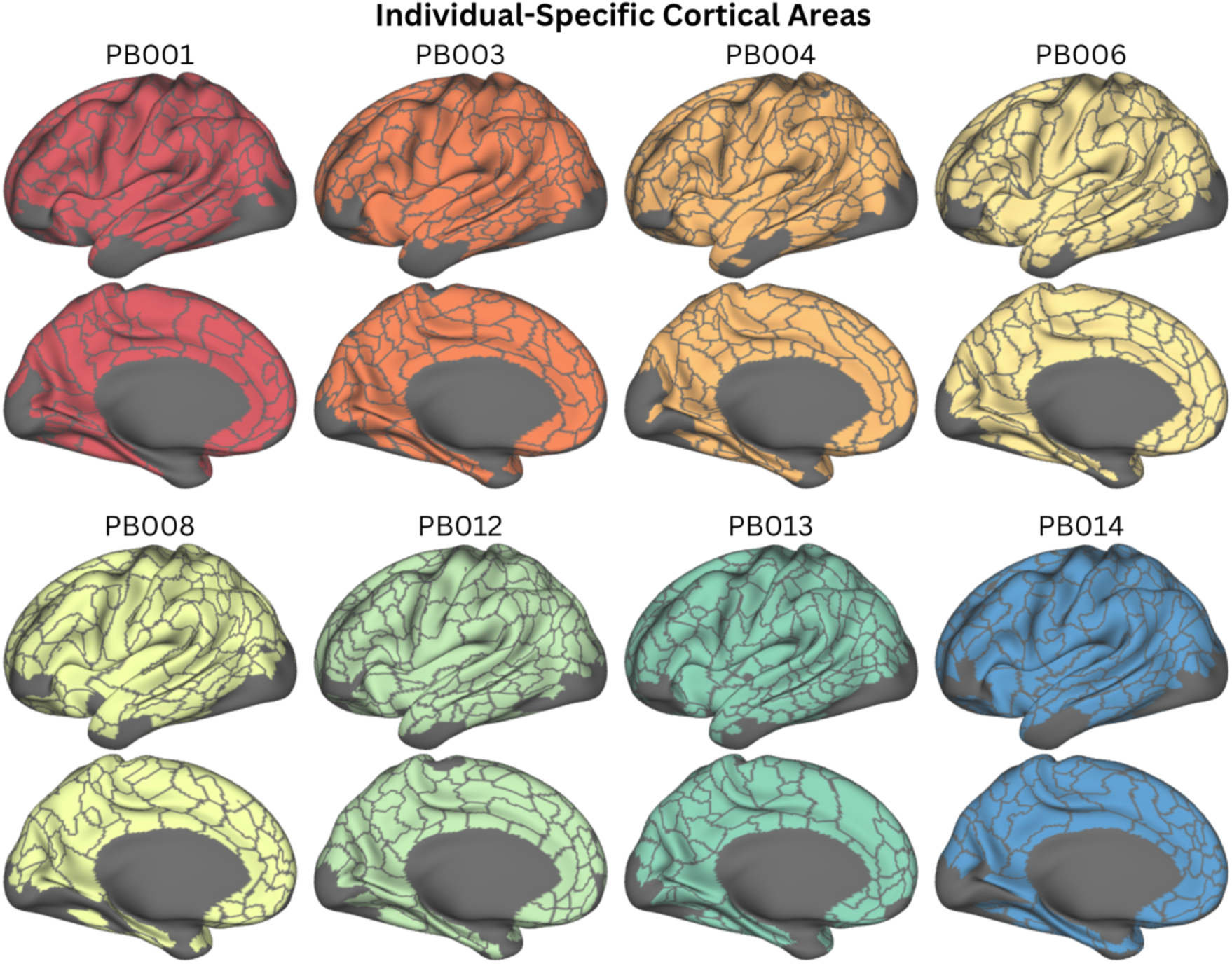
Individual-specific cortical areas. Cortical areas tiling ∼90% of the cortical surface for each individual neonate in the PB dataset projected on the Conte69 Atlas surface (Van Essen et al., 2012). Large patches of the cortex that are unassigned to cortical areas represent areas of low signal-to-noise (SNR) and were excluded from areal parcellations.

**Figure 3.**
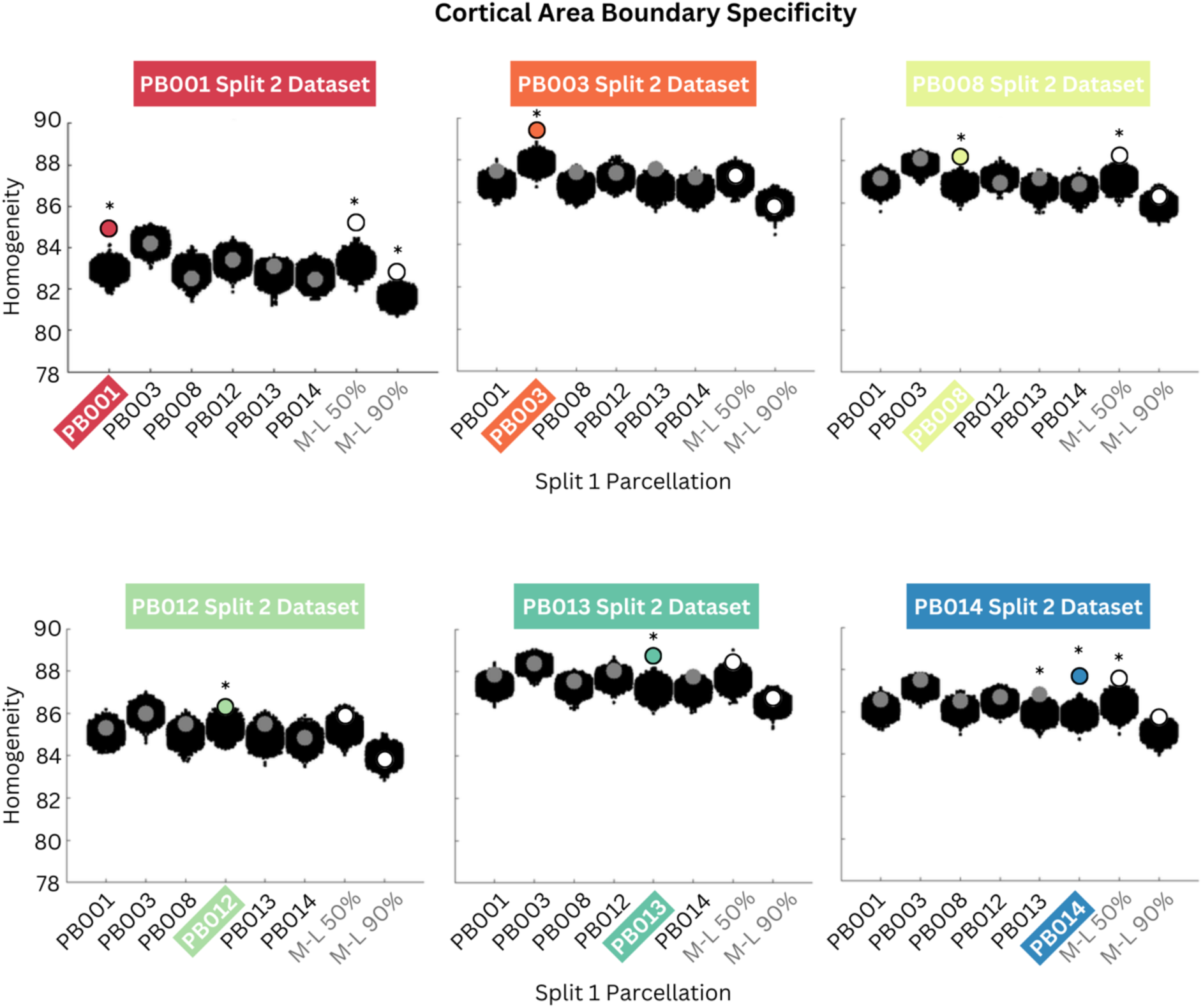
Cortical area boundaries are individual-specific at birth. Data from slit-half 1 were used to generate individual-specific cortical areas for each neonate (x-axis). These individualized parcellations, as well as the ML-50 and ML-90 group parcellations, were evaluated (using homogeneity) against the held-out split-half 2 dataset for each neonate. Large colored dots indicate homogeneity of subjects tested against their own held-out data, large gray dots represent across-subject testing, and large white dots show homogeneity for each subject’s held-out dataset tested against the ML group parcellations. The black clouds of small dots indicate null distributions generated by spatial permutation. Cortical area parcellations that fit significantly better than null parcellations (p < 0.01) are indicated with an *.

### The arrangement of neonatal cortical areas is individually specific

One explanation for why reliable individual neonatal parcellations can cover ∼90% of the cortical surface, but reliable group-average neonatal parcellations can only cover ∼50% of the cortical surface (i.e., parcel centers only) (Myers et al., 2024) is heterogeneity in the arrangement of cortical areas across neonates. To evaluate the individual specificity of cortical area arrangement, we assessed the overall signal homogeneity of cortical parcels in each individual neonate when tested against the held-out data of other individual neonates. In nearly all cases, cortical area parcellations derived from split-half 1 of a neonate did not perform better than chance when tested against any other neonate’s split-half 2 dataset, (**Figure 3**, see position of larger gray dots relative to the smaller black dots depicting homogeneity of spatially permuted null parcellations; **Supplementary Figure 7**). The lowest p-value for the across-subject comparisons is p=0.006 (PB013 areal parcels fit held-out data from PB014); however, the majority of all other across-subject comparisons had p>0.1. The same pattern of results (i.e., individual specificity in cortical area arrangement) was obtained when evaluating areal overlap to assess parcellation fit across subjects (See **Supplementary Figure 8 C & D, Supplementary Results**).

In our previous work, we found that a group-average parcellation covering 90% of the cortical surface (Myers-Labonte-90; M-L 90) was not very reliable. However, a more spatially constrained group-average parcellation covering only ∼50% of the cortical surface (i.e., parcel centers only; Myers-Labonte-50; M-L 50) was highly reliable. We hypothesized that heterogeneity across neonates precluded the M-L 90 from providing a good fit, while the M-L 50 parcels might represent areas of high overlap in the locations of cortical areas across subjects. This hypothesis would predict that the M-L 50 parcels, but not the M-L 90 parcels, would provide reasonable fits to the individual neonatal data. Consistent with this hypothesis, for all neonates, the M-L 50 provided a better fit than the M-L 90 (**Figure 3**; white dots; t=2.97, p=0.03). The M-L 50 provided a fit significantly better than chance (p<0.001) in three of the neonates, while M-L 90 provided a significant fit (p<0.001) in only one neonate (PB001).

The parcellations derived above are based on abrupt transitions in FC across the cortical surface; thus, we evaluated the individual specificity and strength of these abrupt transitions. FC gradients and boundaries, the metrics of abrupt transitions used to derive cortical parcels, were highly reliable within-subject and individually specific (**Supplementary Figure 8 A & B**), consistent with the results above. However, near cortical area boundaries (within 2.5-mm in adult- scaled atlas space), individual neonates exhibited weaker transitions in FC (gradient magnitude) compared to individual adults (t=8.2, p<0.001) (**Supplementary Figure 9**). Additionally, transitions in FC were significantly less abrupt (more gradual slope in gradient magnitude) in neonates compared to adults further away from area boundaries (t=3.5, p=0.02; up to 20-mm in adult-scaled atlas space). These results suggest that, while individually specific near birth, the boundaries between cortical areas are weaker in neonates compared to adults, potentially indicating lower maturity and ongoing development of cortical boundaries in neonates.

### Individual differences in arrangement of neonatal cortical areas are brain-wide

The results above indicate that the overall arrangement of cortical areas in neonates is individually specific, but the possibility remains that the arrangement of *specific* cortical areas in specific locations is more consistent across individuals. For example, the arrangement of cortical areas in early developing sensorimotor regions might be more consistent across individual neonates than in later-developing association regions. To evaluate the across-subject consistency in the locations of specific cortical areas, we generated a whole-brain map that indicates, for each part of the cortical surface, how well cortical areas derived from one neonate fit datasets from other neonates. More specifically, we computed a whole-brain map of the average homogeneity of cortical areas from across-subject comparisons only (**Figure 3A**). We then tested whether this map indicated equal homogeneity across the brain; or if instead across-subject homogeneity was higher in sensorimotor versus association regions.

There was no association between the sensory-association (S-A) axis map (Sydnor et al., 2023) and our whole-brain homogeneity map (r=-0.136, z=-0.807, p=0.419). Additionally, neither the average homogeneity in sensorimotor nor association regions was significantly different than expected by chance based on spatial permutation testing (**Figure 4B**). We next assessed consistency across various adult-defined and infant-defined functional brain networks. Of 35 networks tested, two were significant but did not survive correction for multiple comparisons. (**Supplementary Figure 10**). Across-subject homogeneity in the adult-defined default mode network (DMN) was greater than could be expected due to chance (p<0.05), while across-subject homogeneity in the neonatal-defined medial orbito-frontal network was less than could be expected due to chance (p<0.05). Overall, these results suggest that individual differences in the arrangement of neonatal cortical areas are brain-wide, rather than regionally specific.

**Figure 4.**
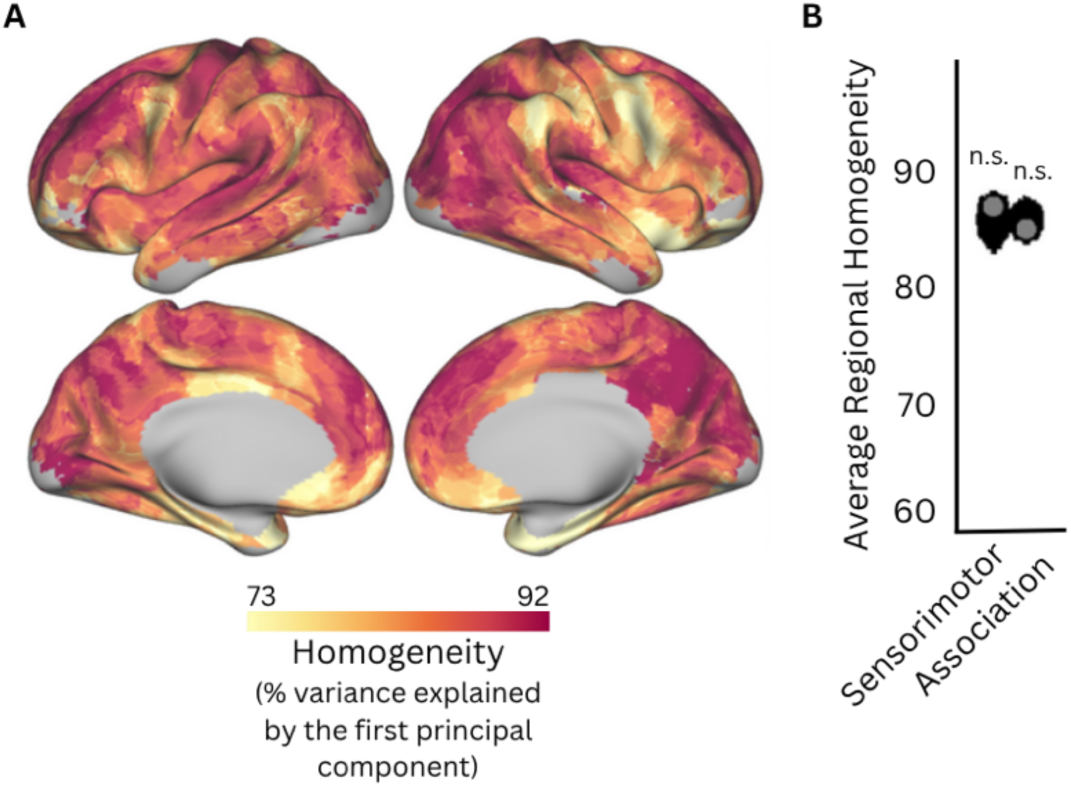
Individual differences in arrangement of neonatal cortical areas are largely brain-wide. **A)** A whole-brain map representing the average homogeneity of cortical areas from across-subject comparisons calculated in Figure 3. **B)** The average homogeneity of cortical areas from across-subject comparisons for sensorimotor and association regions are indicated by large gray dots. The black clouds of small dots indicate null distributions generated by permutation. Neither sensorimotor nor association regions indicated differences in homogeneity greater than chance.

## DISCUSSION

The current study demonstrates the feasibility of collecting large fMRI datasets in neonates, which are necessary to characterize the development of cortical areas in early life. Application of the boundary mapping technique to neonatal PFM data demonstrated that individuals already have reliable cortical areas covering the entire cortical surface near the time of birth. The current study clarified that the prior difficulty in finding reliable cortical areas covering the whole brain using group-average data may be due, in part, to heterogeneity across individuals. Analysis of FC homogeneity between neonates suggested that the consistency of the locations of cortical areas across individual neonates is brain-wide. Altogether, this study demonstrates the maturity of cortical areas near birth and underscores that studies of individuals are required to reveal key aspects of brain development and organization.

### PFM in neonates is feasible and yields reliable and individually specific measures of FC

To accurately characterize functional brain organization in individuals, measures of FC must be reliable. In adults, at least 30-minutes of low motion FC data are needed to measure whole-brain patterns of FC with reliability metrics greater than r=0.85 (Gordon et al., 2017b; Laumann et al., 2015). On average, we observed an average reliability of r=0.78 with the maximum amount of held-out data (41.9-minutes). While we observed increasing reliability with greater amounts of data, we did not observe a plateau in our reliability curves. Thus, it is likely that increasing amounts of data in neonates will yield even greater reliability. Future studies will be required to determine how much data is needed in neonates to reach a plateau in reliability.

Our findings are also consistent with prior work indicating that neonatal FC is less reliable than adult FC. The decreased reliability in neonatal FC data is likely due to a combination of factors including lower connectivity strength in neonates (Arichi et al., 2012; Cusack et al., 2018), varied arousal states during neonatal data collection, and methodological issues. In adults, drowsiness and transitions between sleep states results in lower reliability in FC measures (Tagliazucchi and Laufs, 2014). Neonatal data is collected during natural sleep, and so transitions between sleep states may lower reliability (Mueller et al., 2025; Whitehead, 2025). Neonatal fMRI also has lower signal-to-noise and greater distortions compared to adults (Cusack et al., 2018), which will further decrease reliability. In all, larger amounts of low-motion data (>50-70-minutes) should be collected in sleeping neonates to obtain highly reliable measures of FC comparable to older samples.

### The arrangement of cortical areas is heterogeneous across individuals near birth

Despite showing general consistencies, adults show individual differences in functional brain organization including variation in the location, size, and shape of cortical areas (Gordon et al., 2017b; Laumann et al., 2015). However, it has remained elusive as to when this individual specificity emerges. The neonatal period is an important developmental epoch to investigate the emergence of individual specific functional brain organization due to the rapid and profound increases in FC and structural brain growth happening during this very early postnatal period (Bethlehem et al., 2022; Nielsen et al., 2022). In the current study, the arrangement of cortical areas in any given individual neonate provided a poor fit to the other neonates, suggesting the presence of individually specific functional brain organization already near birth. Possible sources of this individual variability include genetics, influences of the in-utero and postnatal environment, or individual differences in the brain maturity during this time of rapid development.

### Individual differences in the arrangement of neonatal cortical areas are largely brain-wide

Sensorimotor areas demonstrate earlier myelination, FC strength, and functional network maturation compared to higher-order association areas, suggesting cortical development along a sensorimotor to association axis (Doria et al., 2010; Gao et al., 2015a, 2015b, 2009; Larsen et al., 2023; Smyser et al., 2010; Smyser and Neil, 2015). Even still, it remains unknown whether the development of cortical areas in humans follows this same pattern. In animal models, cortical areas develop from a protomap that is refined by thalamocortical inputs postnatally, highlighting the importance of environmental inputs for areal maturation (O’Leary et al., 2007; Petersen et al., 2024). One possibility is that sensorimotor environmental inputs shortly following birth drive the developmental segregation of cortical areas in sensorimotor systems before more complex inputs can drive the developmental segregation of association cortical areas. We hypothesized that the earlier development of sensorimotor areas would be reflected in greater consistency in the arrangement of cortical areas across individuals. However, we did not observe any evidence supporting this hypothesis, and there are at least two possibilities for this result. The first possibility is that consistency in the arrangement of cortical areas across individuals is not related to the maturation of areas.

The second is that cortical area development doesn’t follow the S-A axis trajectory but instead is either homogenous or follows some other pattern. For example, prior work has identified a set of ‘core’ regions within each functional network that are more or less consistent in location across children and adults (Dworetsky et al., 2021; Tu et al., 2025b); one possibility is that areas in these core regions mature prior to other areas. Longitudinal PFM could help adjudicate these possibilities by tracking changes to individual cortical areas over development. Additionally, other functional properties may also lend insight into how cortical areas mature within an individual beginning near birth and how this maturational trajectory relates to individual variation in area arrangement.

### Limitations

Several limitations should be acknowledged in interpreting our findings. As mentioned previously, all data collection during this study occurred during natural sleep. While sleep state and arousal impact the reliability of FC in adults, it is not known how sleep may impact the reliability of FC in neonates. Additionally, it is not known how sleep may impact FC boundary mapping used in the current study to derive cortical areas. Rapid developmental changes near birth complicate determining how much the apparent individual differences relate to slight differences in brain maturity between infants or imaging sessions. Finally, the PB dataset was collected as pilot data and was therefore collected in varying amounts and using various scanning sequences. This heterogeneity in the dataset may complicate across-subject comparisons. However, we don’t expect these differences to impact our findings, as this work focused on within-subject analyses; and we identified significant across-subject variation even among the neonates that had identical scanning paramaters.

### Conclusions

The current study demonstrates feasibility of using PFM to capture reliable individual differences in FC in neonates. Using this technique revealed a fundamental property of neonatal brain organization that is obscured in group-average studies: cortical areas are established near birth, but their arrangement exhibits considerable individual variability. This work opens the exciting prospect of using longitudinal PFM to characterize individualized developmental trajectories and determine how individual differences in these trajectories relate to psychiatric and neurodevelopmental outcomes later in life (Labonte et al., 2024).

## Acknowledgments

We thank the families of all our participants for their invaluable participation and dedication towards this research. We also thank Victoria Brooks and Karen Lukas for their critical roles in participant recruitment, study management, and data collection – your efforts were instrumental to the success of this work. Additionally, we acknowledge the contributions of Joey Scanga and Ramone Agard for their skilled assistance with data curation and preprocessing efforts.

## CRediT Authorship Contribution Statement

**Alyssa Labonte**: Conceptualization, Data curation, Formal analysis, Methodology, Visualization, Writing – original draft, Writing – review and editing; **Julia Moser**: Writing – review and editing; **Cat Camacho**: Writing – review and editing; **Cindy Tu**: Methodology, Writing – review and editing; **Muriah Wheelock**: Writing – review and editing; **Tim Laumann:** Methodology, Writing – review and editing; **Evan Gordon**: Methodology, Writing – review and editing; **Damien Fair**: Conceptualization, Supervision, Writing – Review and editing; **Chad Sylvester**: Conceptualization, Funding acquisition, Supervision, Writing – original draft.

