## Supplementary Materials for "Precision Functional Mapping of the Individual Human Brain Near Birth"

### SUPPLEMENTARY METHODS

*Low Signal Mask for PB Dataset*

The average BOLD images for each run were combined within an individual following mode 1,000 normalization. A single low-signal mask was then generated identifying any vertex with a mean BOLD signal <750. Each individual subject's low-signal mask was combined to form a conjunction map, whereby each vertex was given a value representing the number of individual subjects with low signal at that particular vertex. The final low-signal mask was then generated for any vertex which had low signal for at least half (n=3) of the subjects.

*Cortical Area 'Fit' – DCBC*

To further assess how well the parcels fit the underlying FC data, we used the distance-dependent boundary coefficient (DCBC). The DCBC measures the average difference in whole-brain FC similarity (Pearson's r) between vertices within the same parcel versus those in different parcels, across 1-mm geodesic distance bins. This measure offers an unbiased approach to account for comparing parcellations with a varying number of cortical area parcels across them (Tu et al. 2025; Zhi et al. 2022). The value for DCBC ranges from -1 to 1, whereby the fit of a random parcellation is expected to be zero and a positive DCBC value indicates a fit of the parcellation to the underlying FC data better than chance.

*Cortical Area Spatial Overlap – DSC and ARI*

To quantify the spatial overlap of cortical area boundaries, the Dice similarity coefficient (DSC) was computed on binarized boundary maps (parcels=0, boundaries=1). To assess the significance of the computed DSC, we considered a null distribution of 1,000 randomly rotated parcellations (as described in the methods for homogeneity) each time computing the DSC of the two randomly rotated parcellations to derive a null distribution against which to compare the value obtained in the true parcellations. To quantify the spatial overlap of cortical area parcels, the Adjusted Rand Index (ARI) was computed on non-boundary vertices.

*Gradient Magnitude Evaluation*

To assess the strength of neonatal area boundaries, we quantified the magnitude of the local transitions in FC (i.e., local gradients) and compared their strength to those in identified in individual adults. The individual adult dataset used a point of comparison for this analysis, the Midnight Scan Club, has been previously described (Gordon et al. 2017). For both the neonatal and adult datasets, gradient maps and individual-specific cortical area parcellations were generated for each individual subject (see Parcel Generation in the main text Methods). The average gradient magnitude was calculated for all vertices within a certain distance from their nearest area boundary. Distances from area boundaries were rounded to the nearest 5-mm bin based on their geodesic distance. Bins ranged from 0–25-mm from the nearest cortical area boundary. Average gradient magnitudes for each subject at each distance bin were then averaged to generate plots and compute statistical comparisons between adults and neonates. The slopes as a percentage of the maximum gradient magnitude were calculated as follows:

$$slope\ as\ percentage\ of\ maximum = \left(\frac{slope}{maximum\ gradient\ magnitude}\right) \times\ 100$$

### SUPPLEMENTARY RESULTS

#### Neonatal precision neuroimaging is feasible and is moderately reliable and individually specific.

Measures of FC were moderately reliable within-subject across multiple days, and less similar across-subjects, suggesting individual-specific patterns of FC beginning at birth. Because the amount of low-motion data differed across all split-half datasets, and this may influence across-individual comparisons of both FC reliability and similarity, we repeated analyses of FC reliability and similarity with matching amounts of data across individual neonates. For FC reliability analyses, each held-out split-half contained 31.7 mins of data across all subjects, and data were sampled from all remaining low-motion data for each subject (mean=45.8 mins, range=30-70 mins). FC similarity analyses included equal amounts of data in each split-half across all subjects (31.7 mins). These results are illustrated in **Supplementary Figure 4** and indicate that overall, individual neonates show similarly moderate reliability across scanning sessions (mean (r) = 0.73, SD (r) = 0.05) and are overall less similar to one another than to their own held-out data (within subject: mean (r) = 0.66, SD (r) = 0.05; across subject: mean (r) = 0.52, SD (r) = 0.06; t=2.4, p=0.04).

#### FC gradient-based cortical areas can be reliably defined in individual neonates and cover ~90% of the cortical surface.

Individual-specific cortical area parcellations are reliable across independent data from the same individual. The homogeneity value of each parcel in each individual neonate's cortical area parcellation generated using their full dataset is illustrated in **Supplementary Figure 5A**, and the homogeneity z-score calculated across all cortical area parcels for each individual neonate is illustrated in **Supplementary Figure 5B (**mean z=4.4, all p<0.01). We further assessed the 'fit' of each individual subject's own cortical area parcellation with the underlying patterns of FC using the distance-controlled boundary coefficient (DCBC) (**Supplementary Figure 5C**). Like homogeneity, the DCBC was calculated from cortical area parcellations generated from split-half

1 of each individual subject's dataset and were compared against patterns of RSFC from their own split-half 2 dataset. On average the DCBC for each subject was 0.17 (SEM=0.0199). The average DCBC for each neonate is on the order of what is seen in the adult and older infant literature (Tu et al. 2025; Zhi et al. 2022). These findings suggest that each subject's own cortical area parcellation 'fit' their own unseen data better than a random parcellation.

We further assessed the reliability of individual-specific cortical area parcellations by evaluating the spatial overlap of both cortical area boundaries and the cortical area parcels themselves. To do this, we generated independent cortical area parcellations from each split-half of an individual's dataset and quantified the spatial overlap of their cortical area boundaries and cortical area parcels using the dice similarity coefficient (DSC) and the adjusted rand index (ARI), respectively. Individual maps illustrating the overlap of cortical area boundaries and their respective DSC are shown in **Supplementary Figure 6**. On average, cortical area boundaries generated from independent split-halves of an individual subject's dataset had a DSC of 0.39 (range (DSC)=0.38 – 0.41) which was significantly higher than chance (mean (z) = 9.55, SD (z) = 2.23, $p<0.001$). Spatial overlap of cortical area parcels showed similar significant results with an average ARI of 0.53 (SD (ARI)=0.03), which was also significantly higher than chance.

**The arrangement of neonatal cortical areas is individually specific.**

To evaluate the individual specificity of cortical area arrangement, we assessed the goodness of fit and spatial overlap for cortical parcellations for each individual neonate when tested against the held-out data from other individual neonates. We used the DCBC as an additional metric to determine the 'fit' of individual-specific cortical area parcellations across each neonate. As such, cortical area parcellations were generated from split-half 1 of each neonate's datasets and each resulting parcellation scheme was tested in FC data from split-half 2 across all neonates. **Supplementary Figure 7B** illustrates the DCBC across all within- and across-subject comparisons, whereby higher positive values indicate significantly greater fit than would be expected with a random parcellation.

The individual specificity of cortical area parcellations was also evaluated by spatial overlap of individually specific areal parcellations across each neonate using the DSC and ARI. For all neonates, the ARI and DSC of their own cortical areas and boundaries generated from split-half 1 and spilt-half 2 of their own dataset was higher than the ARI and DSC of their own split-half 1 with split-half 2 of every other neonate (**Supplementary Figure 8C & D**). Such results further suggest individual specificity of neonatal cortical area boundaries.

**SUPPLEMENTARY FIGURES**

| PB ID | Scan Day | Scan Parameters | | | | % Frames Retained | | Minutes Retained | |
|---|---|---|---|---|---|---|---|---|---|
| | | *Multi-Band Factor* | *TE* | *Resolution (x mm)$^3$* | *TR* | | *Mean* | | *Total* |
| PB001 | 1 | ME-5 | 14.2<br>38.93<br>63.66<br>88.39<br>113.12 | 2.0 | 1.76 | 91.0 | 95.0 | 18.4 | 83.0 |
| | 2 | | | | | 99.1 | | 6.7 | |
| | 3 | | | | | 90.7 | | 24.5 | |
| | 4 | | | | | 99.0 | | 33.4 | |
| PB003 | 1 | 4 | 37.0 | 2.4 | 1.2 | 96.3 | 93.7 | 25.9 | 143.8 |
| | 2 | | | | | 93.8 | | 43.9 | |
| | 3 | | | | | 97.6 | | 19.6 | |
| | 4 | | | | | 89.9 | | 36.2 | |
| | 5 | | | | | 90.7 | | 18.2 | |
| PB004 | 1 | 4 | 37.0 | 2.0 | 1.51 | 91.5 | 94.7 | 22.5 | 43.4 |
| | 2 | | | | | 97.9 | | 20.9 | |
| PB006 | 1 | 4 | 37.0 | 2.4 | 1.2 | 88.5 | 91.0 | 11.9 | 36.6 |
| | 2 | | | | | 95.5 | | 12.8 | |
| | 3 | | | | | 89.1 | | 11.9 | |
| PB008 | 1 | 4 | 37.0 | 2.0 | 1.51 | 56.9 | 62.2 | 46.5 | 87.9 |
| | 2 | | | | | 67.5 | | 41.4 | |
| PB012 | 1 | 4 | 37.0 | 2.0 | 1.51 | 63.5 | 69.1 | 38.4 | 72.2 |
| | 2 | | | | | 74.6 | | 33.8 | |
| PB013 | 1 | 4 | 37.0 | 2.0 | 1.51 | 62.4 | 73.6 | 28.3 | 71.9 |
| | 2 | | | | | 65.2 | | 29.5 | |
| | 3 | | | | | 93.2 | | 14.1 | |
| PB014 | 1 | 4 | 37.0 | 2.0 | 1.51 | 61.5 | 76.7 | 27.8 | 84.3 |
| | 2 | | | | | 80.6 | | 36.5 | |
| | 3 | | | | | 88.1 | | 20.0 | |

**Supplementary Table 1.** *fMRI Scanning Parameters for the Precision Baby (PB) Dataset.* All data for a single participant were collected with identical scanning parameters, though scanning parameters differed across participants. Percent of frames retained included only runs that were included in analyses and not runs that were completely excluded from analyses. PB001 was collected using a Multi-Echo (ME) sequence with 5 echoes and an in-place acceleration factor (IPAT=2).

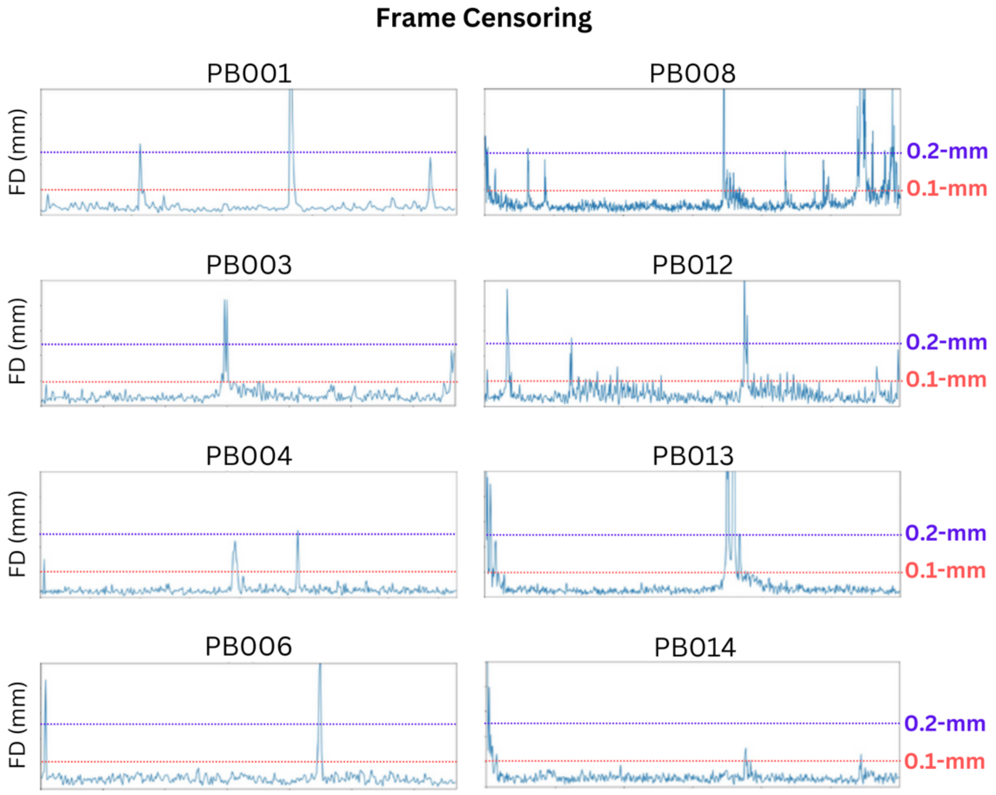


**Supplementary Figure 1.** *Examples of BOLD frame censoring using frame displacement (FD) traces for each individual neonate.* Traditional resting-state fMRI analyses utilize an FD threshold of 0.2-mm. This threshold is too liberal for this particular dataset; therefore, an FD threshold of 0.1-mm was utilized for the PB dataset.

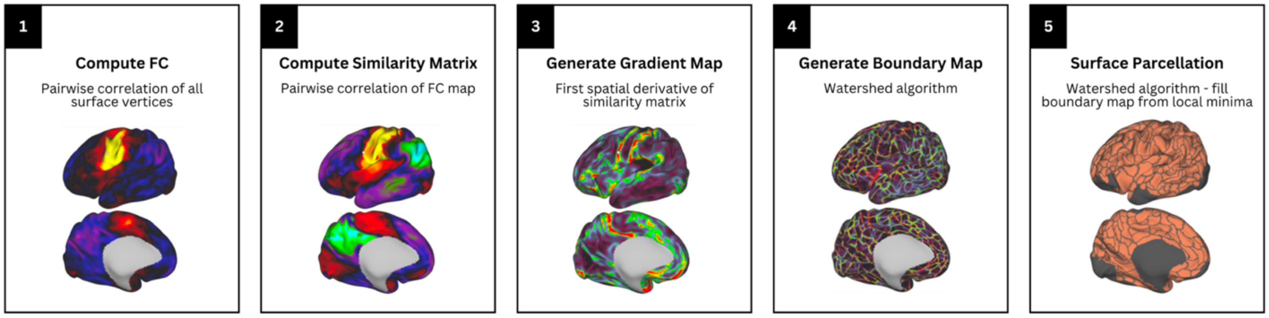


**Supplementary Figure 2.** *Flowchart of cortical area parcellation generation methods*. (1) Whole-brain FC maps at each cortical surface vertex were computed. (2) FC map similarity between all vertices was computed. (3) Spatial gradients were calculated on each column of this similarity matrix using Workbench tools (wb_command -cifti-gradient), creating a gradient map for each surface vertex. (4) Edges were identified in each smoothed gradient map using the watershed edge detection algorithm. These edge maps are averaged across all vertices, creating an edge density map. (5) Parcels were identified by applying the watershed edge detection algorithm to the edge density map. Figures in steps 1-3 represent a single map from a vertex-wise computation.

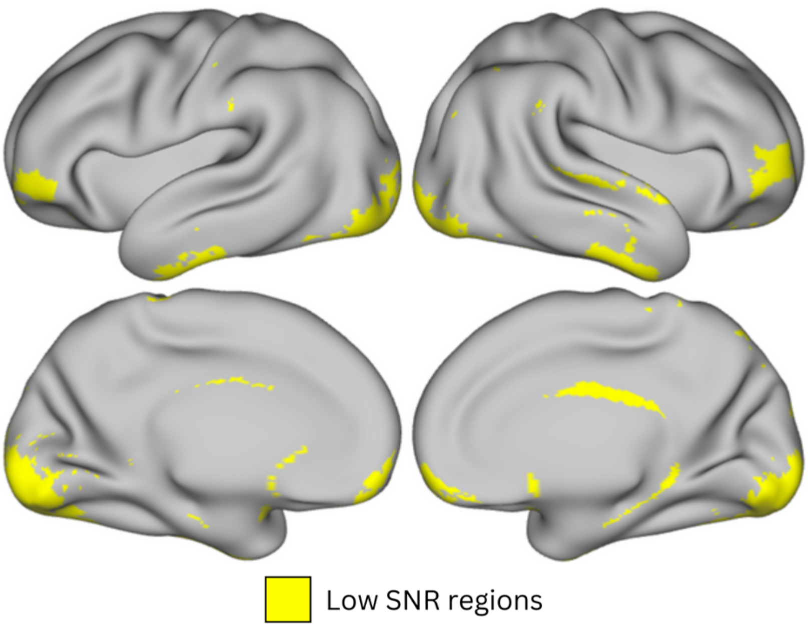


**Supplementary Figure 3.** *Neonatal low SNR mask.* Vertices with BOLD signal < 750 across at least half of all PB participants were included in the Low SNR mask. Parcels with at least 15 vertices inside of any low SNR region (yellow areas) were excluded from the final parcellation and all parcel evaluation procedures.

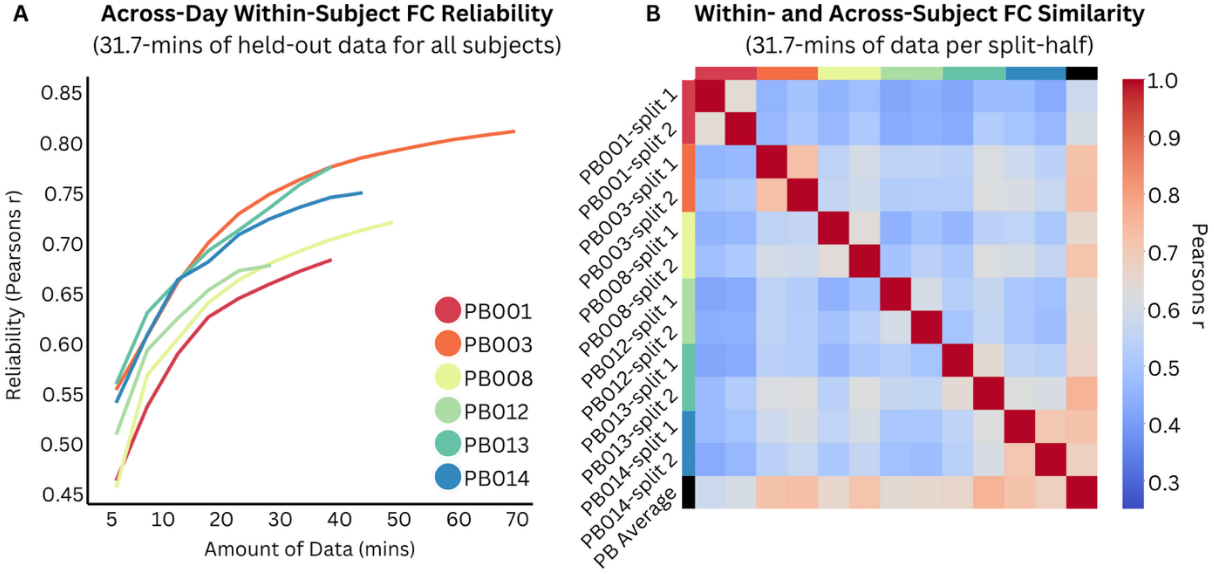


**Supplementary Figure 4.** *Brain-wide patterns of RSFC are reliable and individual-specific at birth using equal amounts of low motion data across all individual subjects.* **A)** Across-day reliability of whole-brain RSFC increases with data quantity. Held-out data for each neonate consisted of 31.7 minutes of low motion RSFC data. Reliability was measured as follows: For each participant, the upper triangle of the FC matrix of a given amount of randomly selected data was correlated with the upper triangle of the FC matrix from the held-out half of their data (31.7-mins) to obtain a Pearsons r correlation for that particular amount of data. This procedure was repeated 1000 times for each participant. **B)** Pairwise similarity of FC matrices between all individual split-halves of data across all participants, as well as the group average (black; last row and column). Each split-half of data across all neonates consisted of 31.7 minutes of low motion RSFC data. Similarity was measured as follows: For each participant, the upper triangle of the FC matrix for each split-half was correlated with the upper triangle of every other split-half FC matrix across all participants, as well as the group average FC matrix.

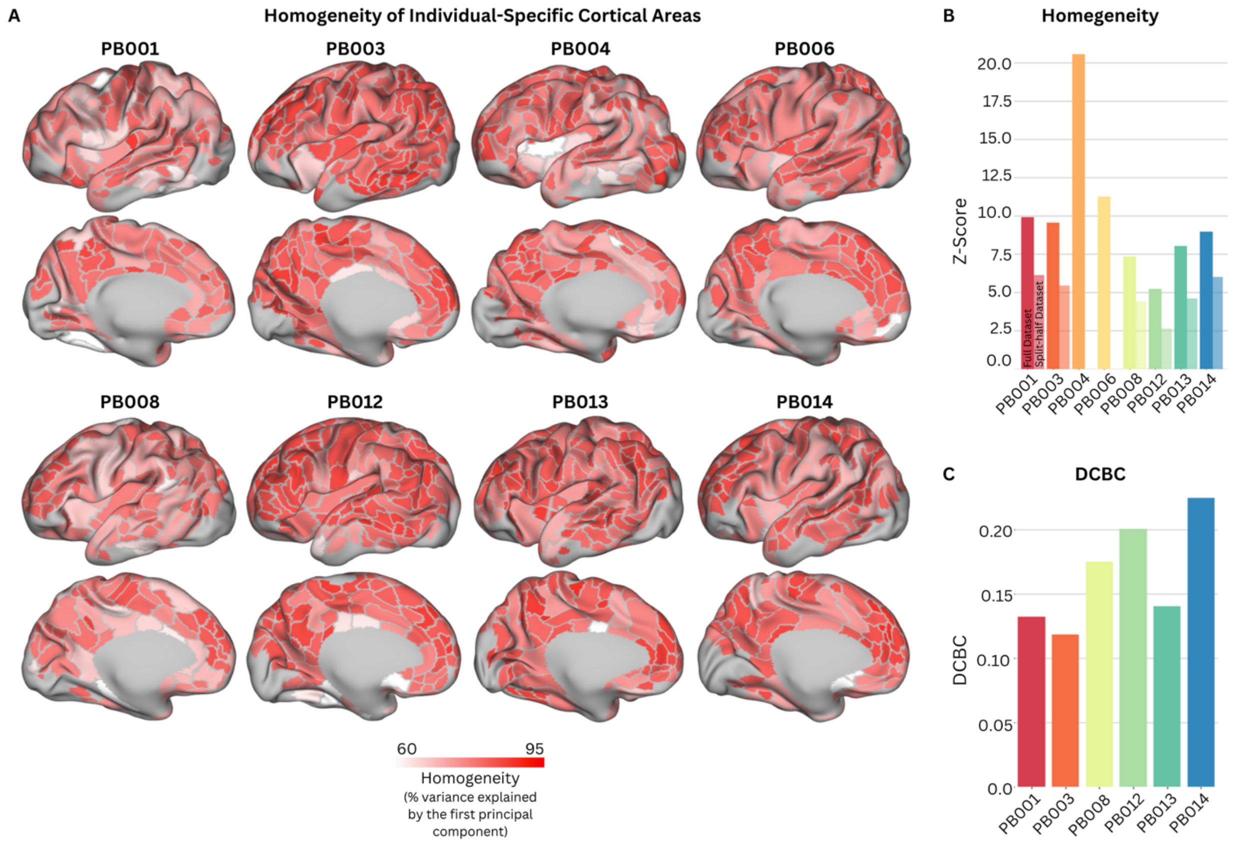


**Supplementary Figure 5.** *Individual-specific cortical areas fit their own RSFC data better than a random cortical area parcellation.* **A)** Homogeneity values for each parcel are shown across all individual's based on cortical areas generated from their own full dataset tested in their own full dataset. Homogeneity is defined as the percent of the variance in RSFC patterns explained by the parcel's first PCA eigenvariate. **B)** Homogeneity was calculated using their individual-specific cortical areas generated from their full dataset on their full dataset (Full dataset, left-most bar for each participant). Additionally, for each participant, homogeneity was calculated using their individual-specific cortical areas generated from split-half 1 of their dataset on their split-half 2 dataset (split-half dataset, right-most bar for each participant). Null model parcels are created by randomly rotating the true parcels x1000 - maintaining each parcel's size, shape, and relative position to one another. The average homogeneity across all cortical area parcels was compared to the average homogeneity across all cortical area parcels from the 1000 null models to calculate a Z-Score. **C)** The DCBC for each neonate was calculated using a subject's own individual-specific cortical areas generated from split-half 1 of their dataset on RSFC data from split-half 2 of their own dataset. Values greater than zero indicate a fit better than chance.

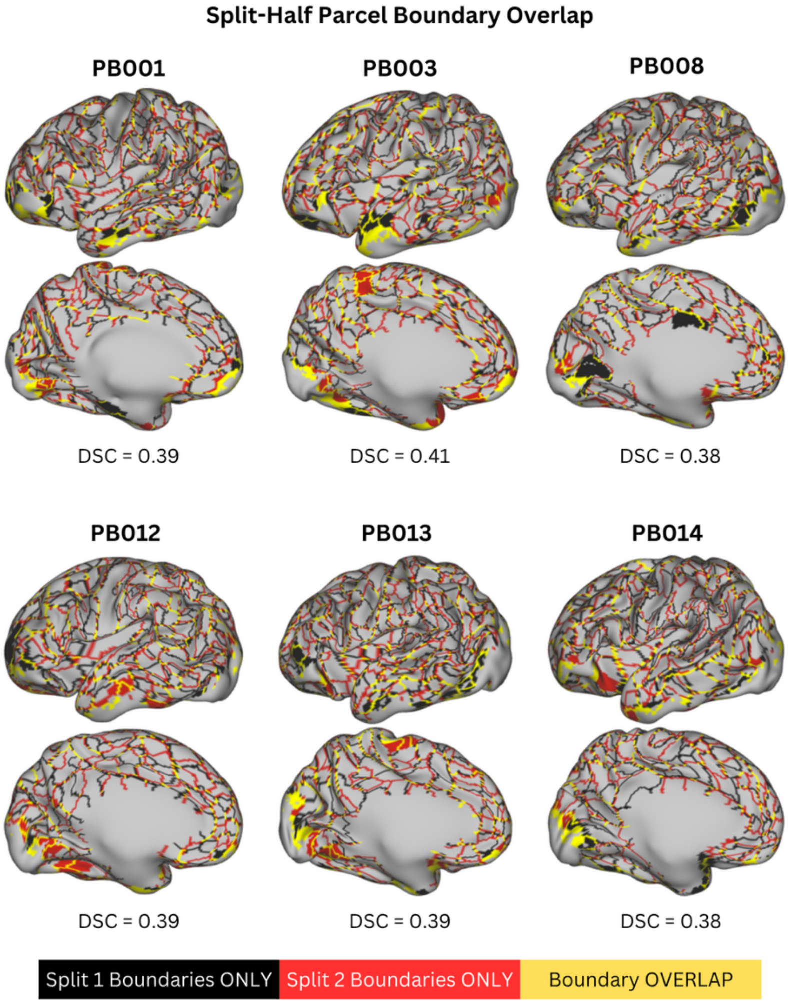


**Supplementary Figure 6.** *Cortical areas boundaries are reliable within an individual neonate.* Data from each split-half of an individual participant's dataset were used to generate individual-specific cortical area boundaries for each neonate. Identified boundaries were overlaid on one another and the Dice Similarity Coefficient (DSC) was calculated on the binary boundary map for each subject. Boundaries identified only in split-half 1 of the dataset are shown in black and boundaries identified in only split-half 2 of the dataset are shown in red. The locations in which the boundaries identified in both split-half 1 and split-half 2 occupy the same vertices are shown in yellow.

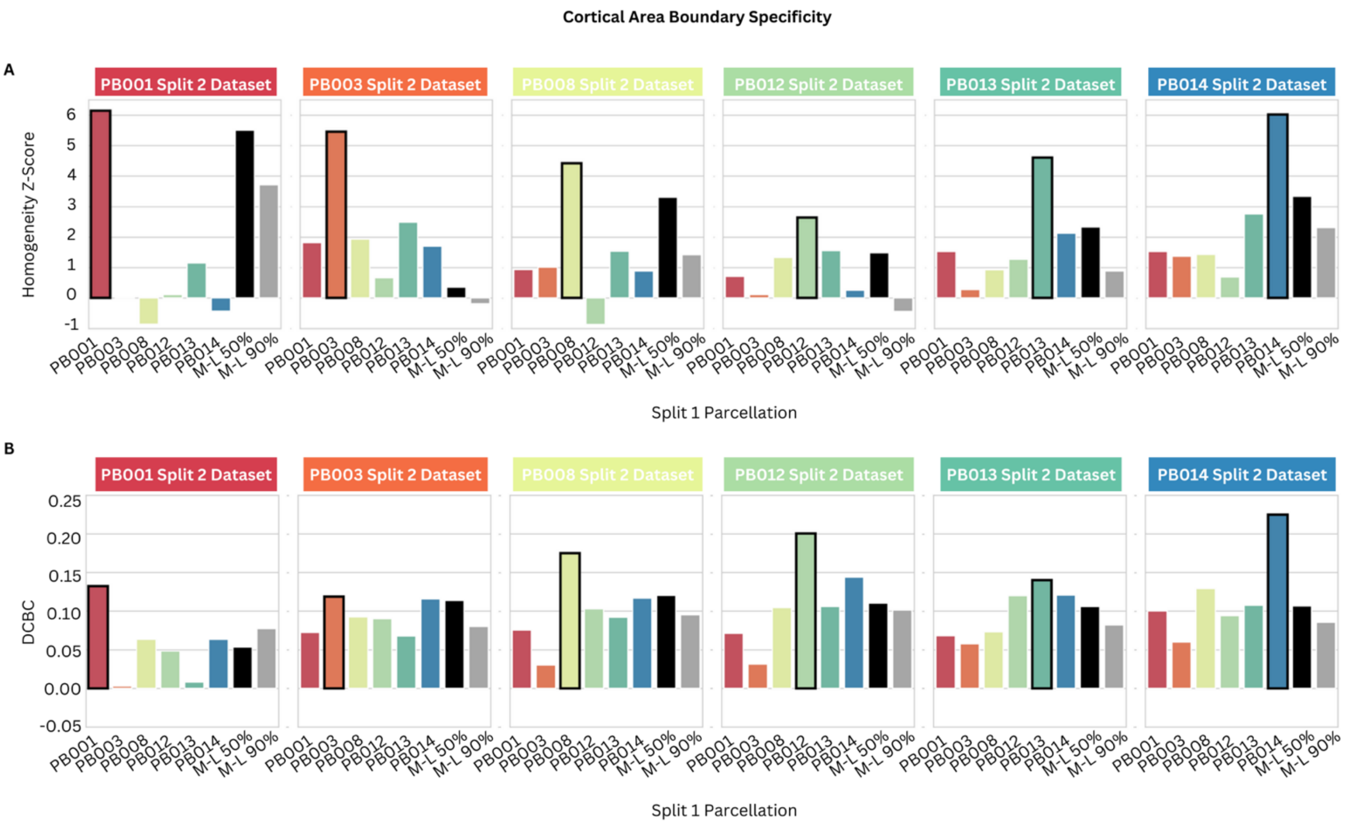


**Supplementary Figure 7.** *Cortical area boundaries are individual-specific at birth.* Data from split-half 1 were used to generate individual-specific cortical area parcellations for each neonate. These parcels were evaluated (using homogeneity **(A)** and DCBC **(B)**) in RSFC data from split-half 2 within and across all neonates, as well as the Myers-Labonte neonatal group parcellation (y-axis).

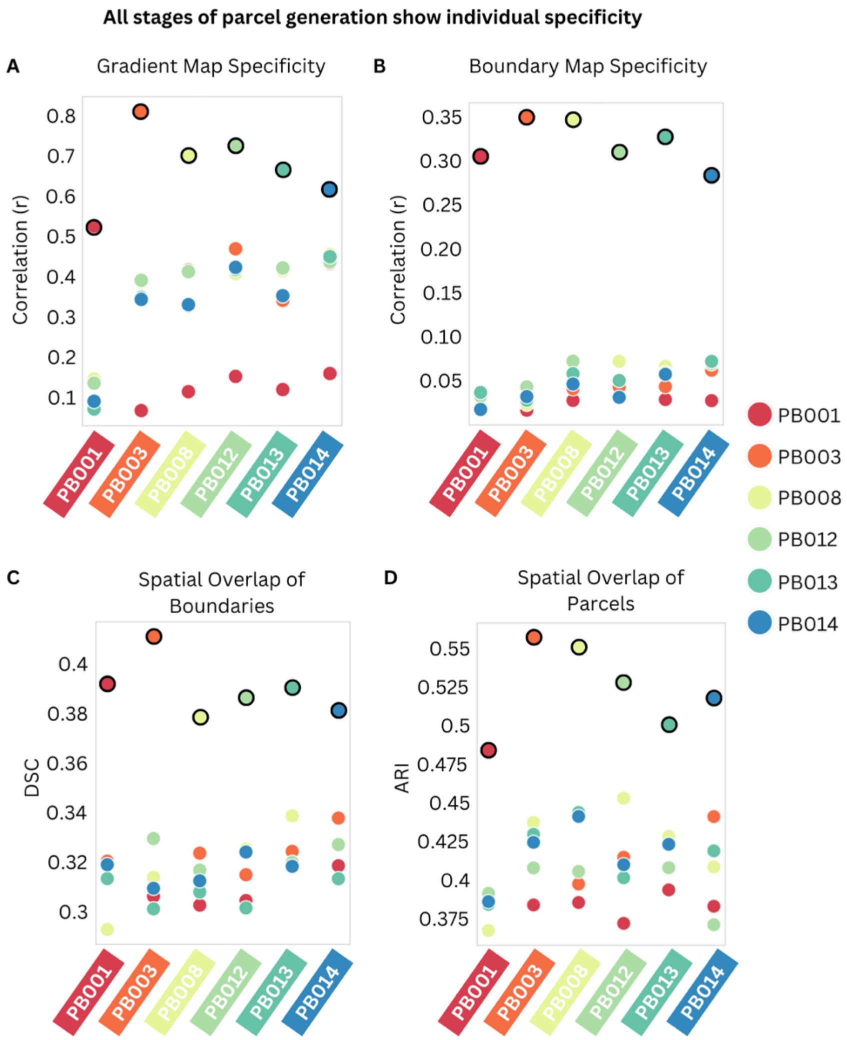


**Supplementary Figure 8.** *All stages of cortical area parcellation generation indicate individual-specific cortical area boundaries at birth.* Data from each split-half of an individual subject's dataset were used to generate individual-specific cortical area parcellations. Individual-specificity was evaluated at each intermediate stage in the process. **A)** Individual-specific gradient maps generated from split-half 1 of each participant's dataset were correlated with gradient maps generated from split-half 2 within and across all neonates. **B)** Individual-specific boundary maps generated from split-half 1 of each participant's dataset were correlated with boundary maps generated from split-half 2 within and across all neonates. **C)** The spatial overlap of individual-specific cortical area boundaries generated from split-halves 1 and 2 was calculated using the DSC within and across all neonates. **D)** The spatial overlap of individual-specific cortical area parcels generated from split-halves 1 and 2 was calculated using the ARI within and across all neonates.

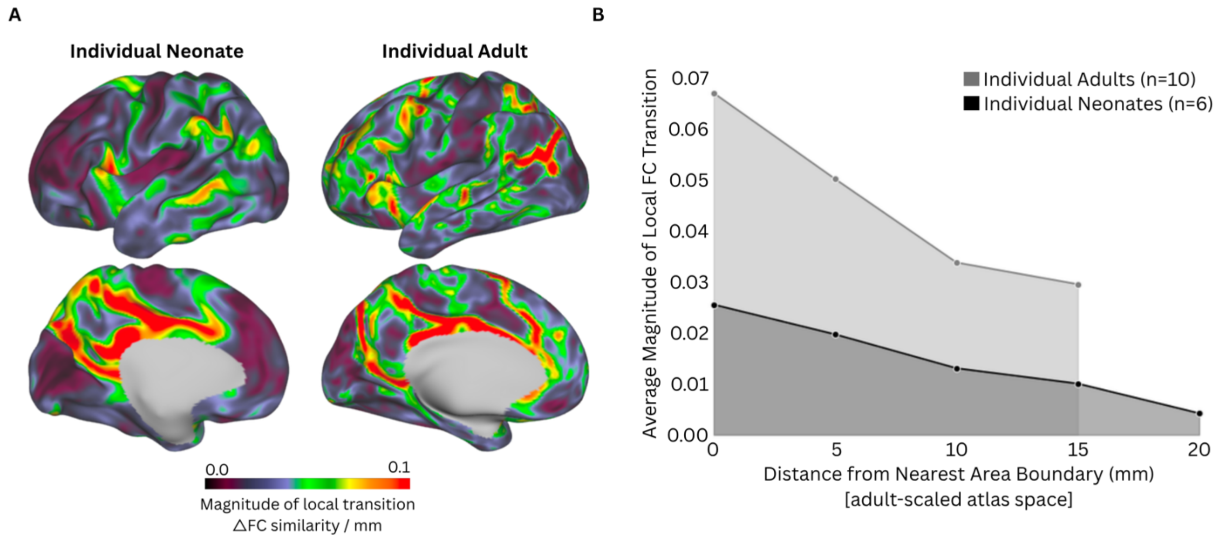


**Supplementary Figure 9.** *The strength of local transitions in FC in individual neonates is weaker than individual adults.* **A)** Local FC gradient representing the magnitudes of transitions in FC between adjacent FC patterns in an individual neonate (left) and in an individual adult (right). **B)** Strength of the of the average local transition magnitude at varying distances (in adult-scaled atlas space) from areal boundaries. Individual neonates exhibited overall weaker transitions in FC compared to individual adults (t=8.2, p<0.001). Individual neonates also exhibited more gradual transitions in FC further from area boundaries compared to individual adults (neonatal slope [as a percentage of maximum gradient magnitude] = -3.8%, adult slope [as a percentage of maximum gradient magnitude] = -4.1%, t=3.5, p=0.02).

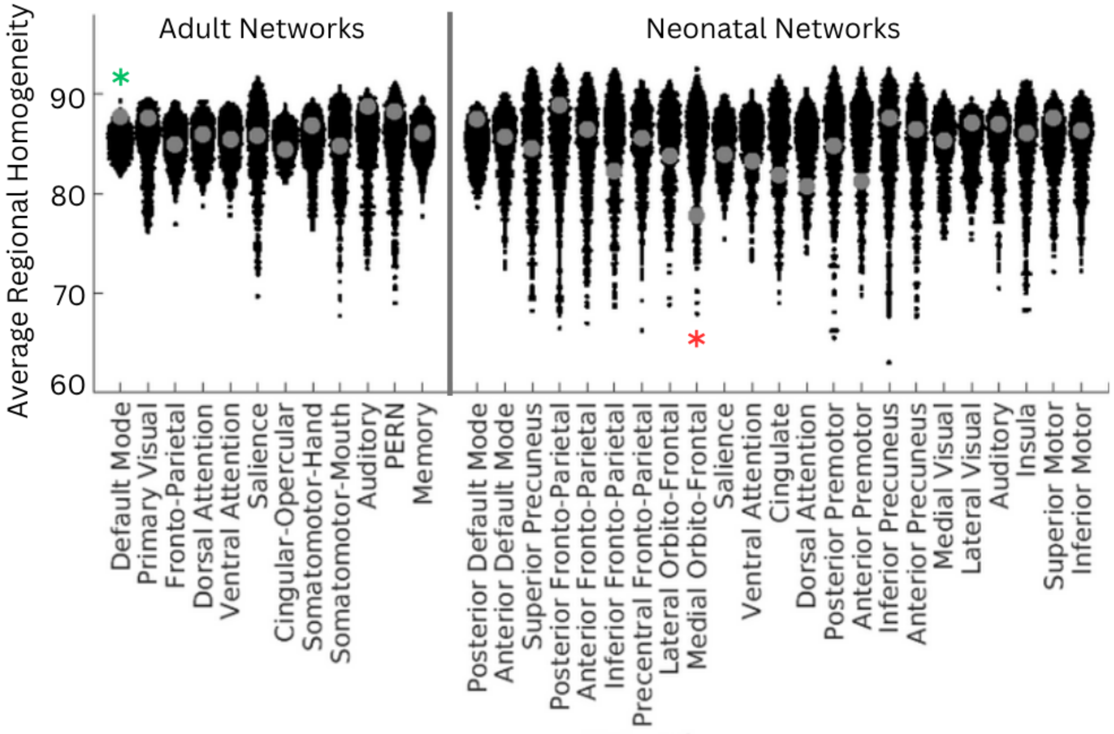


**Supplementary Figure 10.** The average homogeneity of cortical areas from across-subject comparisons for each adult (left) and neonatal (right) functional network are indicated by large gray dots. The black clouds of small dots indicate null distributions generated by permutation. Brain regions or networks which had significantly greater homogeneity than spatially permuted null parcellations ($p < 0.05$) are indicated with a green asterisk, and those which had significantly lower homogeneity than spatially permuted null parcellations ($p < 0.05$) are indicated with a red asterisk. Results are not multiple comparisons corrected across the tests for each network.
